# In vitro investigation of antibacterial activity of Gum Arabic prepared by two different processing methods against Enterococcus faecalis

**DOI:** 10.1101/2023.08.16.553607

**Authors:** Nuha Elmubarak, Yahia Ibrahim, Abbas Gareeballah, Nada Sanhouri

## Abstract

**Background:** Entrococcus faecalis is a known cause of endodontic treatment failure. Synthetic drugs have been preferred for decades, but recently, many plants have been reported for their antibacterial activity.

**Aim:** To investigate the antibacterial effect of Gum Arabic (GA) processed with two different processing methods against Enterococcus faecalis.

**Method:** Antibacterial susceptibility tests against Enterococcus faecalis (ATCC 29212) were performed for 200mg/ml ethanolic extracts of spray-dried and mechanically ground GA using Agar disc diffusion. Sodium Hypochlorite (1%), Chlorhexidine (0.2%), and Antibiotic multi-disc were used as positive controls, and ethanol (20%) as a negative. The inhibition zones diameters were measured.

Serial dilutions of both types of Gum Arabic (100, 50, 25, 12.5 mg/ml) were tested for their antibacterial activity.

**Results:** In Concentration 200 mg/ml, spray**-**dried GA displayed a significantly greater inhibition zone against E. faecalis than mechanically ground *(P=0.02).* Both types of Gum Arabic exhibited lower antibacterial activity than chlorhexidine (0.2%). However, only mechanically ground GA showed significant result *(P=0.005).* Spray-dried GA showed significantly higher antibacterial activity against than Tetracycline 300mcg *(P=0.005)*.

The antibacterial activity of spray-dried GA exceeded that of mechanically ground in all concentrations of serial dilutions, except for 12.5mg/ml, both are similar.

**Conclusion:** processing method of Gum Arabic affects its antibacterial potency against E. faecalis. In high concentrations, spray-dried GA is active antibacterial, while mechanically ground is non-active.

Decreasing the concentration of mechanically ground GA increases its inhibitory effect, but the opposite effect was observed with spray-dried GA.

## Introduction

Enterococcus faecalis is a gram-positive, facultatively anaerobic coccus bacteria that can survive harsh conditions (1, 2). It is part of the habitual flora in the mouth, human gastrointestinal tract, and female genital tract. E. faecalis is a known cause of endodontic treatment failure and some systemic diseases. It has developed high resistance to antimicrobial agents (1).

Isolates from oral infections differ from hospital-derived isolates as they do not present many mobile genetic elements. However, oral isolates have virulence factors related to adhesion and biofilm formation, which help the bacteria to colonize different oral sites. Moreover, oral strains may also carry specific antibiotic resistance determinants that have the potential to be transferred to other pathogenic bacteria in biofilm communities (2).

For many years, synthetic drugs were commonly used. However, nowadays, herbal medicines are gaining popularity. In fact, there have been studies on various plants and their potential to prevent the growth of microorganisms (3). These medicinal plants are continuously being screened for their effectiveness. One of the ten most promising medicinal plants in Africa is Gum Arabic or Acacia Senegal (4). This gum is naturally produced from Acacia trees (5). Sudan is the leading producer of Acacia Gums worldwide, followed by Nigeria, Chad, Mali, and Senegal (6).

Acacia senegal was screened for phytochemicals and found to contain tannins, glycosides, and flavonoids, as well as oxidases, peroxidases, and pectinases, all of which have antimicrobial properties (7–10).

Many studies have proved the antibacterial efficacy of Gum Arabic against various bacterial species (7, 8, 11–15).

A considerable number of medicinal plants have been investigated for their antimicrobial potency against Enterococcus faecalis (16). No studies consider the effect of Gum Arabic against Enterococcus faecalis.

Gum Arabic, undergoes different processing methods depending on the desired quality. There are two common methods: mechanical milling and spray drying. In mechanical milling, dried gum nodules are crushed to create a granular powder. On the other hand, in spray drying, a solution of Gum Arabic is pasteurized and then sprayed into hot air to evaporate all the water and leave a dry powder of Gum Arabic. The latter method has an advantage over raw gum as it is free from microbial contamination and dissolves more quickly (10).

The aim of this study is to investigate the antibacterial effect of Gum Arabic processed by mechanical grinding and spray-drying techniques against Enterococcus faecalis.

## Material and method

### Collection of the Plant

Gum nodules were collected and authenticated at the Herbarium of Medicinal and Aromatic Plants & Traditional Medicine Research Institute with code number G-1983-1-MAPTRI-H. After that, nodules were dried on air at room temperature away from the sun and processed in two different processing methods:

1. Spray drying technique.
2. Manual grinding with a pestle and mortar.

Three Gum Arabic samples were sent to the National Public Health Laboratory to ensure that they met microbiological standards. These samples included the spray-dried GA, the manually ground GA (with a pestle and mortar), and ready-made mechanically ground powder from a private company.

The analysis showed that the ready-made mechanically ground sample was contaminated with E-coli. Moreover, Yeasts and Molds have also been detected with levels exceeding the acceptable limits in this sample. The other two samples were matched with the Microbiological Standard for Gum Arabic. However, the spray-dried GA showed better results than the manually ground sample. These two samples were the samples used in this study.

### Preparation of extract

For each type of Gum Arabic, three samples of 50-gram weight were extracted with ethanol of different concentrations - 70%, 80%, and 99.9%. The samples were soaked in different concentrations of ethanol (Reagents Duksan, Ethyl Alcohol, absolute, product No.6923, 2.5L)and then left for three days while being filtered daily. Afterward, the solvents were evaporated, and the extracts were freeze-dried until they were completely dry. The yield percentages of the three ethanolic extracts (70%, 80%, 99.9%) of both types of Gum Arabic were compared.

### Antibacterial susceptibility test

Antibacterial susceptibility tests were conducted for the 70% ethanolic extract of both types of Gum Arabic against Enterococcus faecalis using the Agar disc diffusion method as per Elkhateeb 2017 (17), with the following steps:

#### 1. Preparation of the media

Muller-Hinton Agar (HIMEDIA, REF. M173-500g Mueller Hinton Agar) was prepared according to the instructions. Each 20 mL of the freshly prepared and autoclaved Muller-Hinton agar was poured into a sterile petri dish and maintained at room temperature to cool down. Before use, the plates were incubated at 35 °C for 48 hours, and the sterility was checked.

#### 2. Preparation of discs

Filter paper discs of 6 mm diameter were prepared from Whatman filter paper No. 1, placed in a petri dish, and sterilized in a hot air oven at 160 °C for two hours.

#### 3. Preparation of extracts solutions

The extract from each type of Gum (200mg) was dissolved in 1ml of 20% ethanol (Reagents Duksan, Ethyl Alcohol, absolute, product No.6923, 2.5L). The mixture was mixed well using a vortex to ensure complete dissolution.

To avoid synergistic effect of the solvent (ethanol 20%) on the antibacterial activity of the extract:

1. Checking the inhibition zone of ethanol 20%: A disc impregnated with ethanol 20% was placed on a plate inoculated with E. faecalis and incubated at 37°C for 24 hours under anaerobic conditions. The solvent was used after no inhibition zone was observed around the disc.
2. Selection of the negative control: Ethanol 20% itself served as a negative control.

#### 4. Reactivation and inoculation of Enterococcus faecalis

To reactivate the standard strain Enterococcus faecalis (ATCC 29212), it was placed in 5ml of brain heart infusion (BHI) broth (HIMEDIA, REF. M210-500G Brain Heart Infusion Broth) and kept under anaerobic conditions for 48 hours. Next, a sterile cotton swab was used to transfer E. faecalis from the BHI broth onto a Muller Hinton Agar plate. The plate was then incubated at 37 degrees Celsius under anaerobic conditions for 24 hours.

#### 5. Preparation and standardization of inoculum suspension

Three to five well-isolated colonies of the same morphological type were selected from a Muller Hinton Agar plate culture and transferred with a sterile loop into a tube containing 5 ml of BHI broth. The broth was incubated anaerobically at 37 °C for 24 hours.

The turbidity of E. faecalis suspension was adjusted equal to that of 0.5 McFarland standard (1.5 × 10^8^ CFU/ml) by using BHI broth for dilution of E. faecalis suspension.

#### 6. Antibacterial susceptibility test

Plates inoculated with Enterococcus faecalis were prepared by streaking the surface of Muller Hilton Agar plates with the adjusted bacterial suspension using a sterile cotton swab.

Three sterile discs were immersed in 10 µL of both extract solutions of Spray-dried Gum Arabic and mechanically ground Gum Arabic. After saturating the discs, they were moved to prepared plates.

Discs were also immersed in Ethanol (20%) to be used as a negative control, while Chlorhexidine 0.2% (Clenora mouthwash, Chlorhexidine Gluconate BP 0.2%w/v) as well as Sodium hypochlorite 1% (Prevest Denpro Hyposol, 3% Sodium hypochlorite 500 ml) to be used as positive controls transferred to the prepared plates.

Ready-made antibiotic multidisc for Gram-positive isolates (Axiom laboratories, Code No. 001, Mantola Pahar Ganj-New Delhi-110055) was transferred to the prepared plate. The antibiotics included in the multidisc for Gram-positive were displayed in (Figure 1).

**Figure 1:**
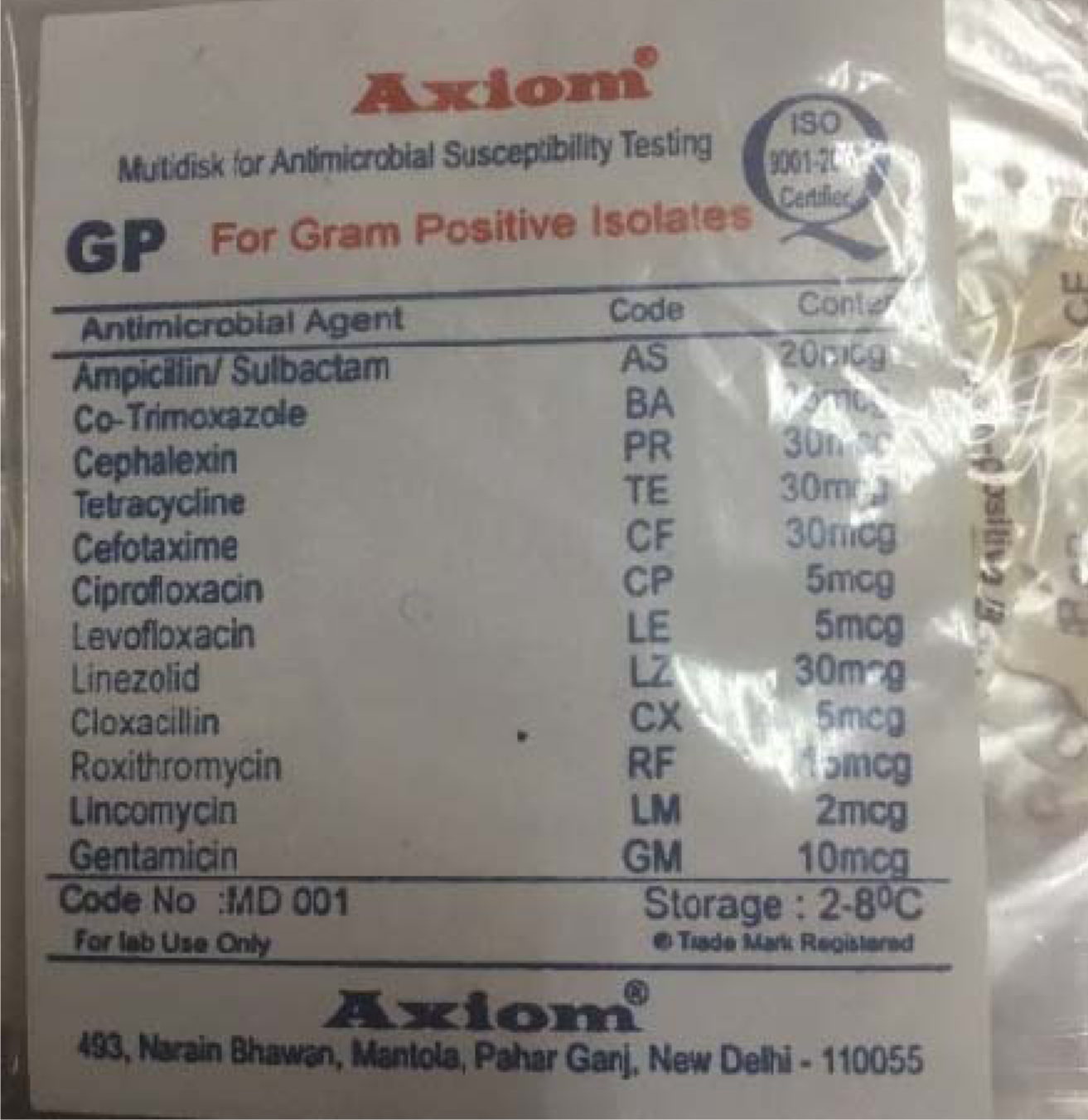
List of antibiotics in multi-disc for antibacterial susceptibility test to gram positive bacteria.

These plates were incubated at 37°C for 24 hours under anaerobic conditions. The plates were observed for the emergence of a clear zone around each disc (inhibition zone). The diameters of the inhibition zones were recorded.

#### 7. Antibacterial susceptibility test for serial dilutions of Gum Arabic (100, 50, 25, and 12.5 mg/ml) against Enterococcus faecalis

Both types of Gum Arabic were coded and diluted in four serial dilutions to obtain concentrations of 100, 50, 25, and 12.5 mg/ml.

Two Muller-Hinton Agar plates were prepared, and labeled according to the extract codes. Each plate was then divided into four labeled zones indicating the extract concentrations. Sterile cotton swab was rolled in the Enterococcus faecalis suspension to streak the surface of Muller Hinton Agar.

For both types of Gum Arabic, four sterile discs were immersed in 10 µL of each concentration (100, 50, 25, 12.5 mg/ml). The discs were transferred to their labeled zones in the coded plates. The plates were incubated at 37°C for 24 hours under an anaerobic condition, which was generated by a CO2 generating kit (Thermo Scientific, AnaeroJar™ 2.5L with Oxoid CO2 Gen Sachet). After incubation, the diameter of the inhibition zones were measured.

An excel plot to display the relation between the concentration of Gum Arabic and the inhibitory effect was conducted for each type of Gum Arabic. The square of the radii diameter of the inhibition zone was presented on the Y-axis, and the log concentrations of Gum Arabic on the X-axis. A suitable curve was drawn from the plot for each type of Gum Arabic. Simple regression analysis was performed to determine the relation between the concentration of Gum Arabic and its inhibitory effect as well as the concentration of each type of Gum Arabic which shows no inhibition (18).

Radius inhibition zone equal ½ [Diameter of inhibition zone in millimeter including disc diameter _ disc diameter] (18)

## Results

### The amounts of ethanolic extracts obtained from spray dried and mechanically ground Gum Arabic

Despite using the same amount of Gum Arabic (GA) and ethanol concentration, the amount of ethanolic extract gained by spray-drying and mechanical grinding techniques showed different results.

The 70 % ethanolic extract of spray-dried GA was more than double that of mechanically ground GA of the same weight. Similarly, 80% ethanolic extract of spray-dried GA was more than triple that of mechanically ground GA extract of the same weight. However, 99.9% ethanol produced no extract from spray-dried GA, while it produced 0.01 mg from mechanically ground GA. Tables 1 provides more details on these findings.

**Table 1:** Yield percentages of both types of Gum Arabic extracted by three different concentrations of ethanol.

| Concentration<br>of ethanol used<br>for extraction | weight of<br>Gum<br>Arabic<br>(Grams) | Spray dried Gum Arabic |  | Mechanically ground Gum Arabic |  |
| --- | --- | --- | --- | --- | --- |
|  |  | weight of extract<br>(Grams) | Yield % | Weight of extract<br>(Grams) | Yield % |
| 70% | 50 g | 0.0815 g | 0.163% | 0.0374 g | 0.075% |
| 80% | 50 g | 0.1011 g | 0.202% | 0.0326 g | 0.065% |
| 99.9% | 50 g | No extract (zero) | zero | 0.006 g | 0.012% |

A thin layer chromatography test (TLC) confirmed that 99.9% ethanol did not produce any extract from spray-dried Gum Arabic.

### Antimicrobial susceptibility test against E. faecalis

The two types of Gum Arabic inhibited the growth of E. faecalis with variant degrees. (Table 2) (Fig. 2-7).

**Table 2:** Measurements of the inhibition zones diameters of Gum Arabic against E. faecalis.

| Investigated material | Inhibition zone diameter (mm) |  |  |  |  |  |
| --- | --- | --- | --- | --- | --- | --- |
|  | M1 | M 2 | M3 | N | mean | Std. dev. |
| Spray dried Gum Arabic (200 mg/ml) | 14 | 15 | 15 | 3 | 14.67 | 0.58 |
| Mechanically ground Gum Arabic (200 mg/ml) | 9 | 8 | 9 | 3 | 8.67 | 0.58 |
| Ethanol 20% (-ve) | 0 | 0 | 0 | 3 | 0 | 0.00 |
| Sodium hypochlorite (1%) (+ve) | 12 | 12 | 12 | 3 | 12 | 0.00 |
| Chlorhexidine (0.2%)(+ve) | 15 | 16 | 17 | 3 | 16 | 1.00 |
| CP (Ciprofloxacin 5mcg) | 20 |  |  | 1 |  |  |
| LE (Levofloxacin 5mcg) | 20 |  |  | 1 |  |  |
| BA (Co-Trimoxazole 25 mcg) | 15 |  |  | 1 |  |  |
| TE (Tetracycline 300mcg) | 10 |  |  | 1 |  |  |

(M1), (M2), and (M3): measurements of inhibition zones around first, second and third discs
and (N): number of measurements for each extract

**Figure 2:**
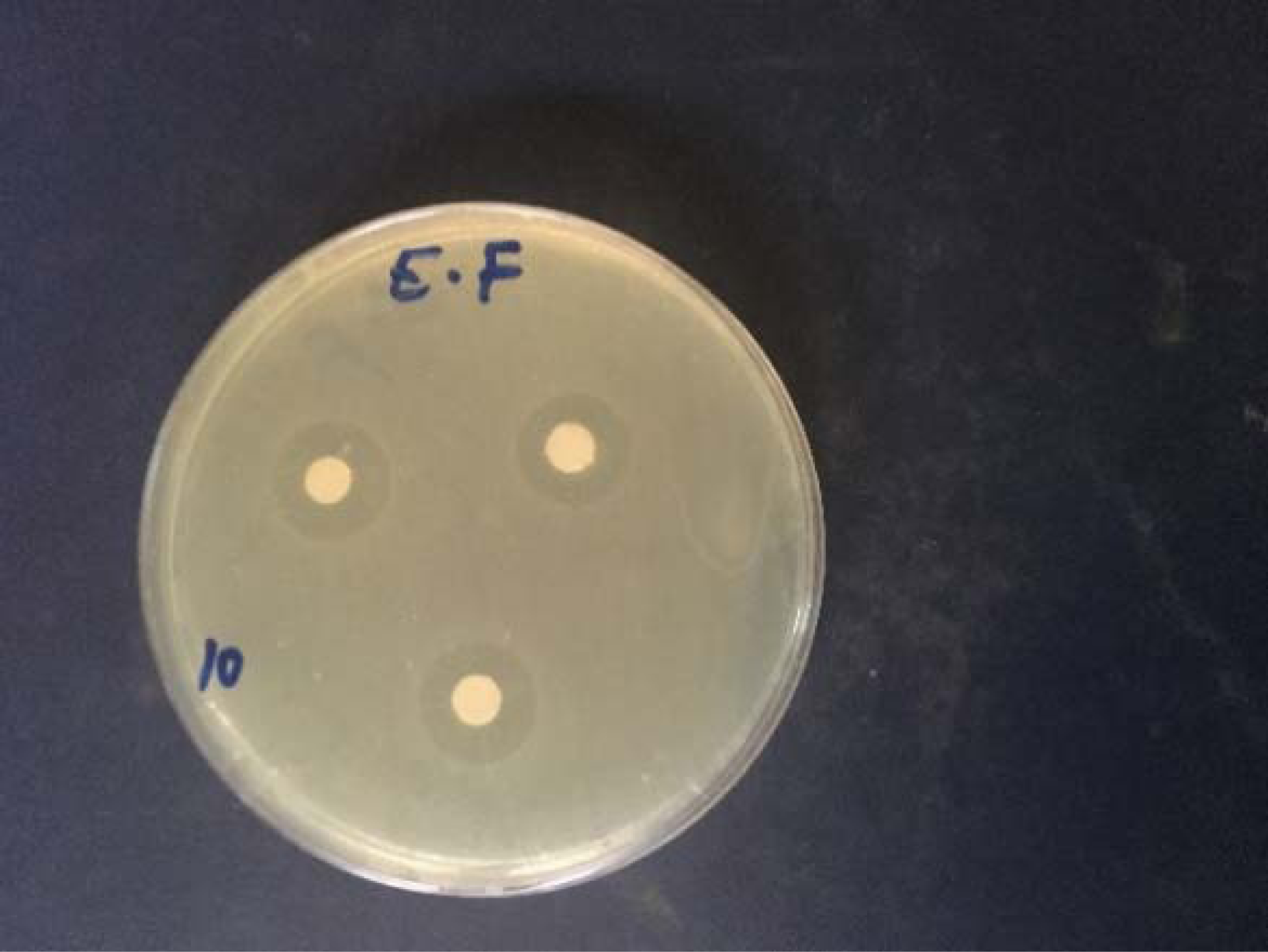
Inhibition zones around discs saturated with spray dried Gum Arabic on a plate inoculated with E. faecalis.

**Figure 3:**
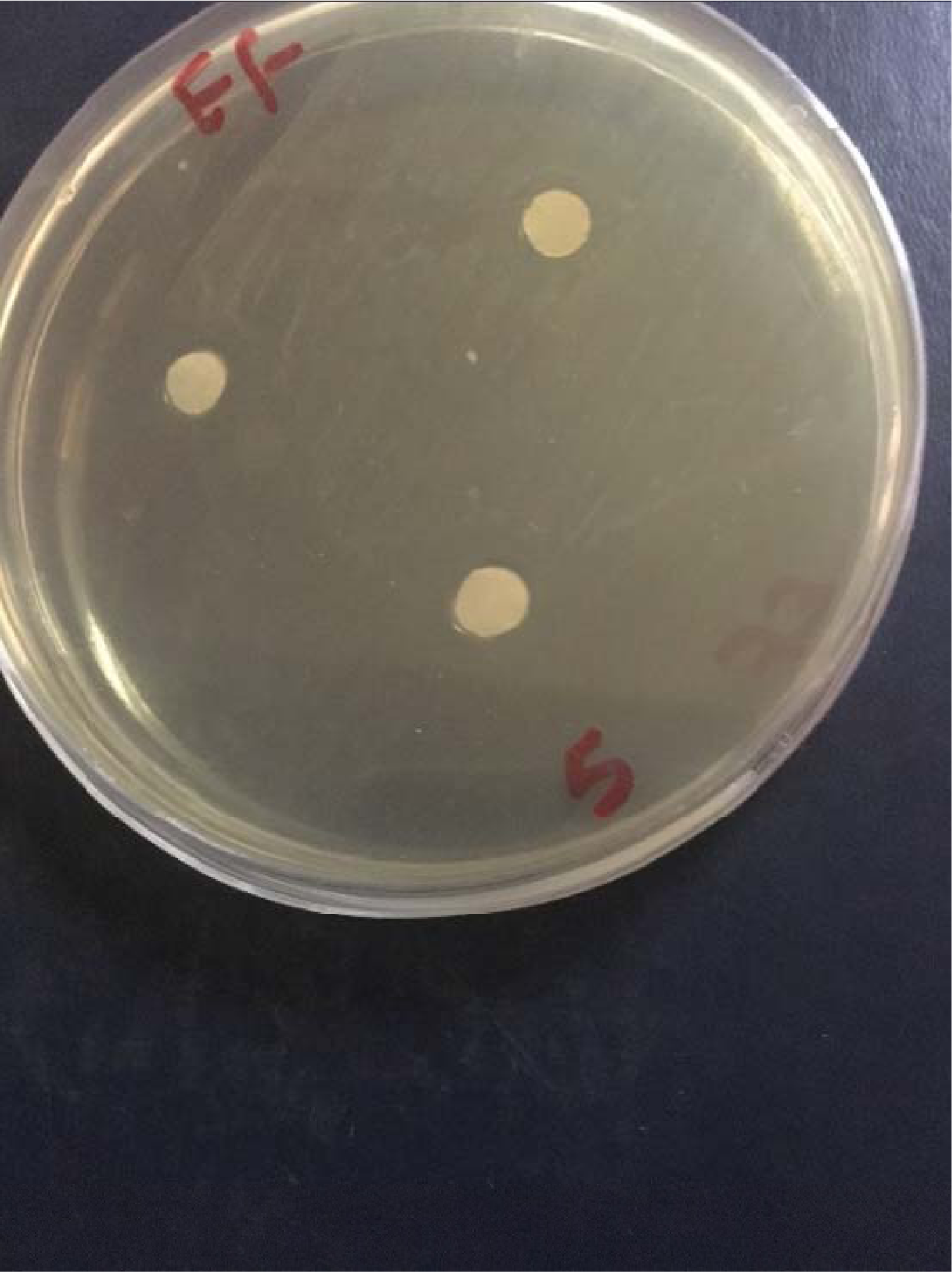
Inhibition zones around discs saturated with mechanically ground Gum Arabic on a plate inoculated with E. faecalis.

**Figure 4:**
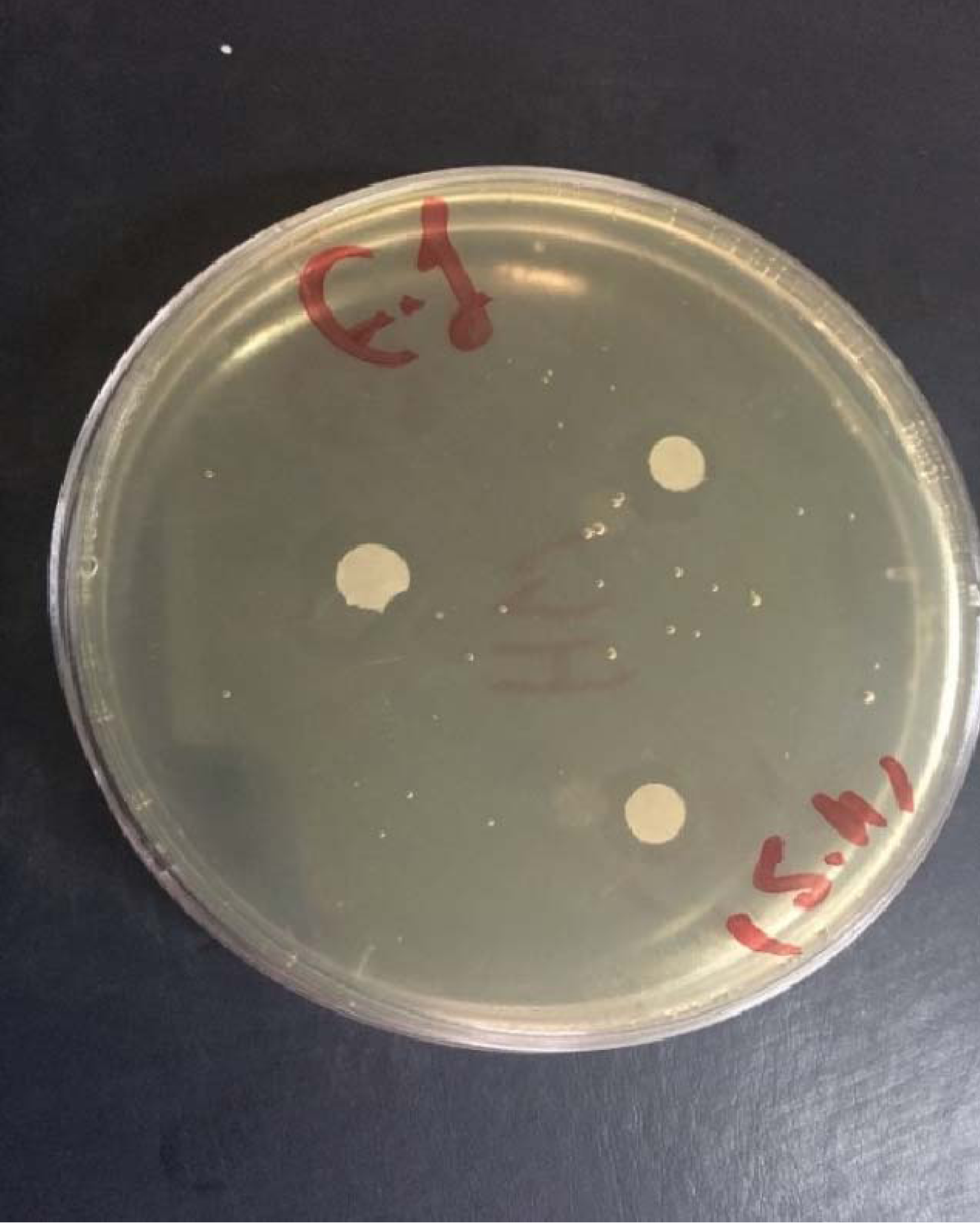
Inhibition zones around a discs saturated with Sodium hypochlorite 1% on a plate inoculated with E. faecalis.

**Figure 5:**
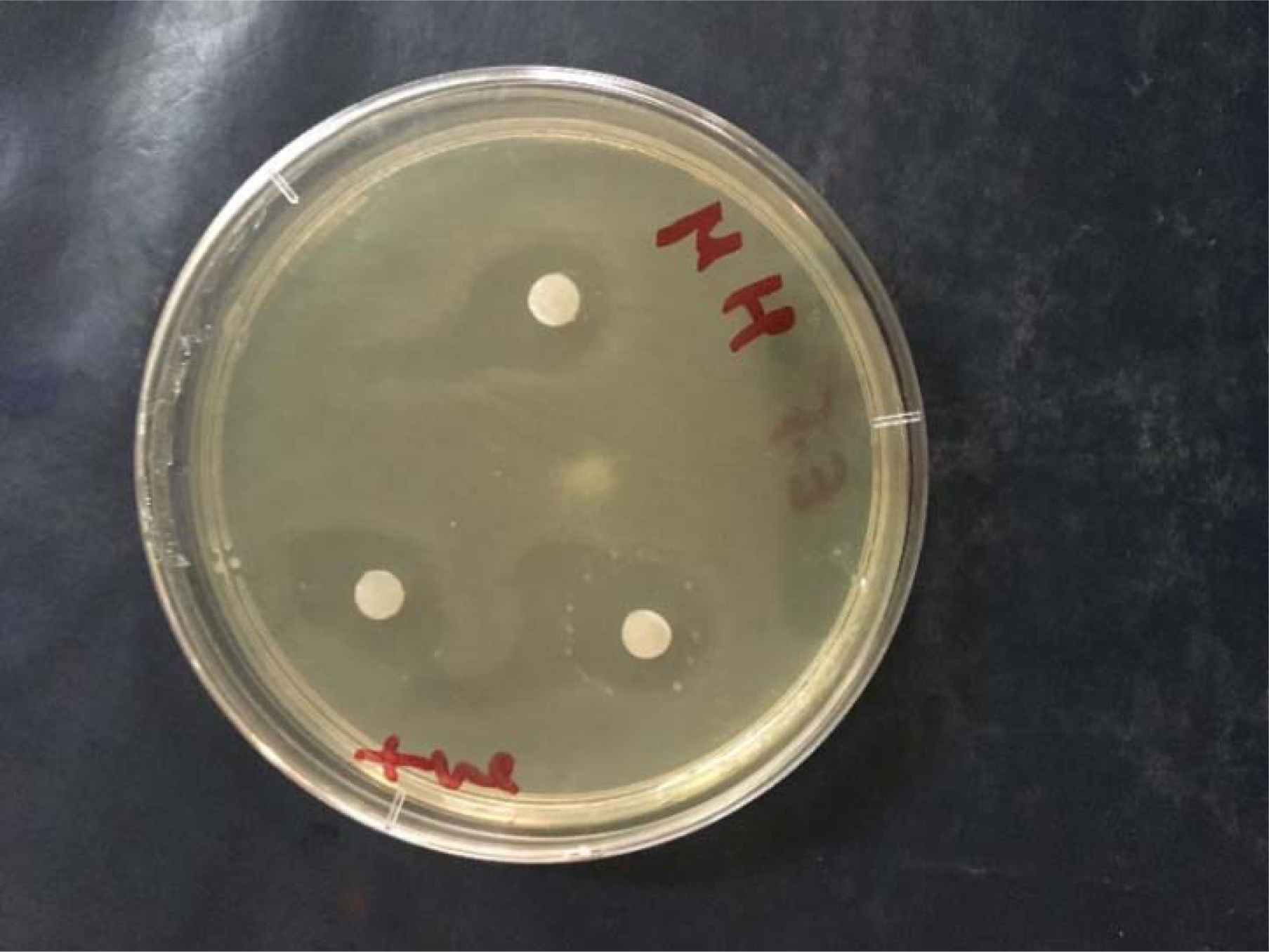
Inhibition zones around discs saturated with Chlorhexidine 0.2% on a plate inoculated with E. faecalis.

**Figure 6:**
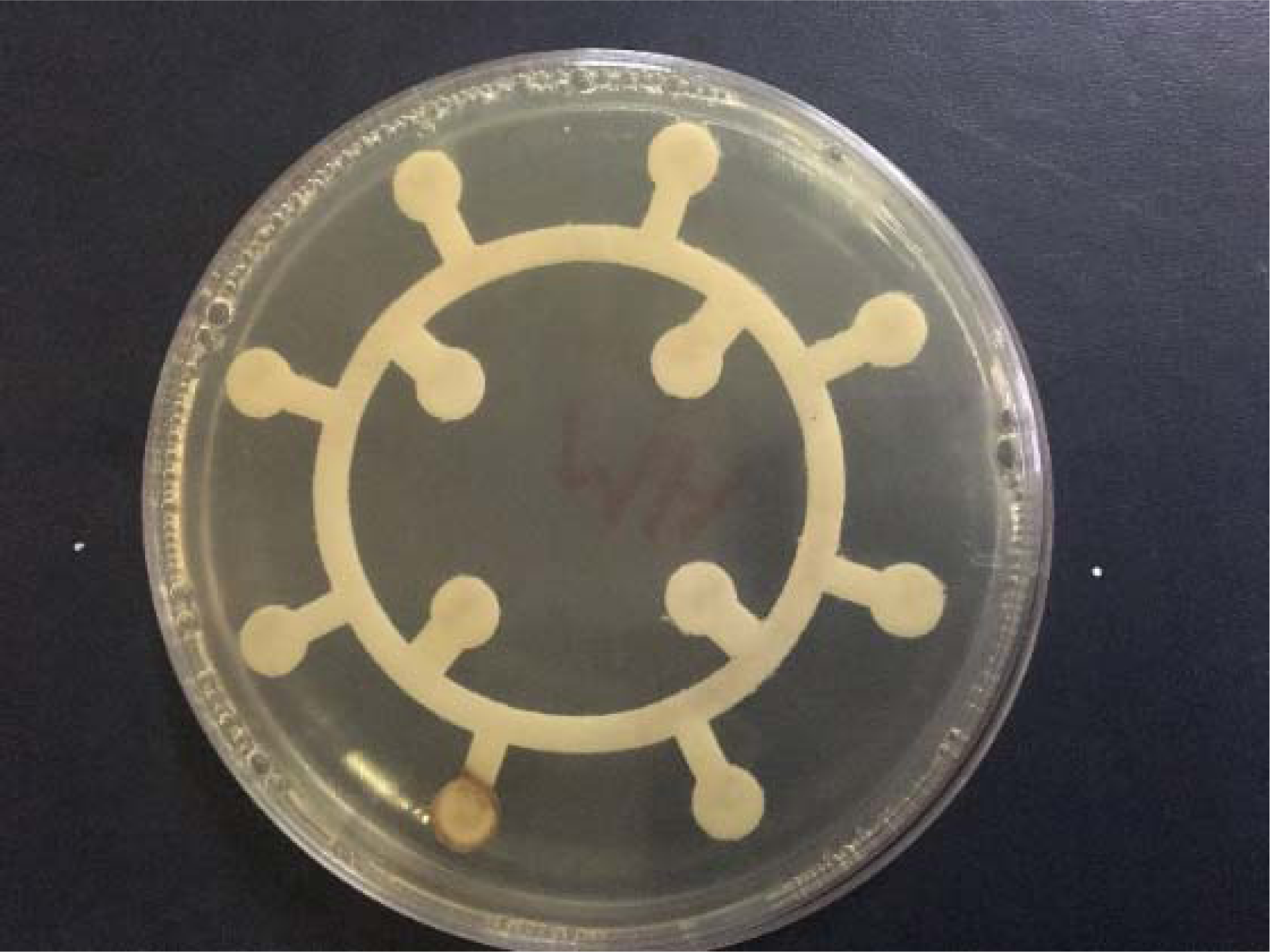
Inhibition zones in multidisc antibiotic for Gram positive bacteria on a plate inoculated with E. faecalis.

**Figure 7:**
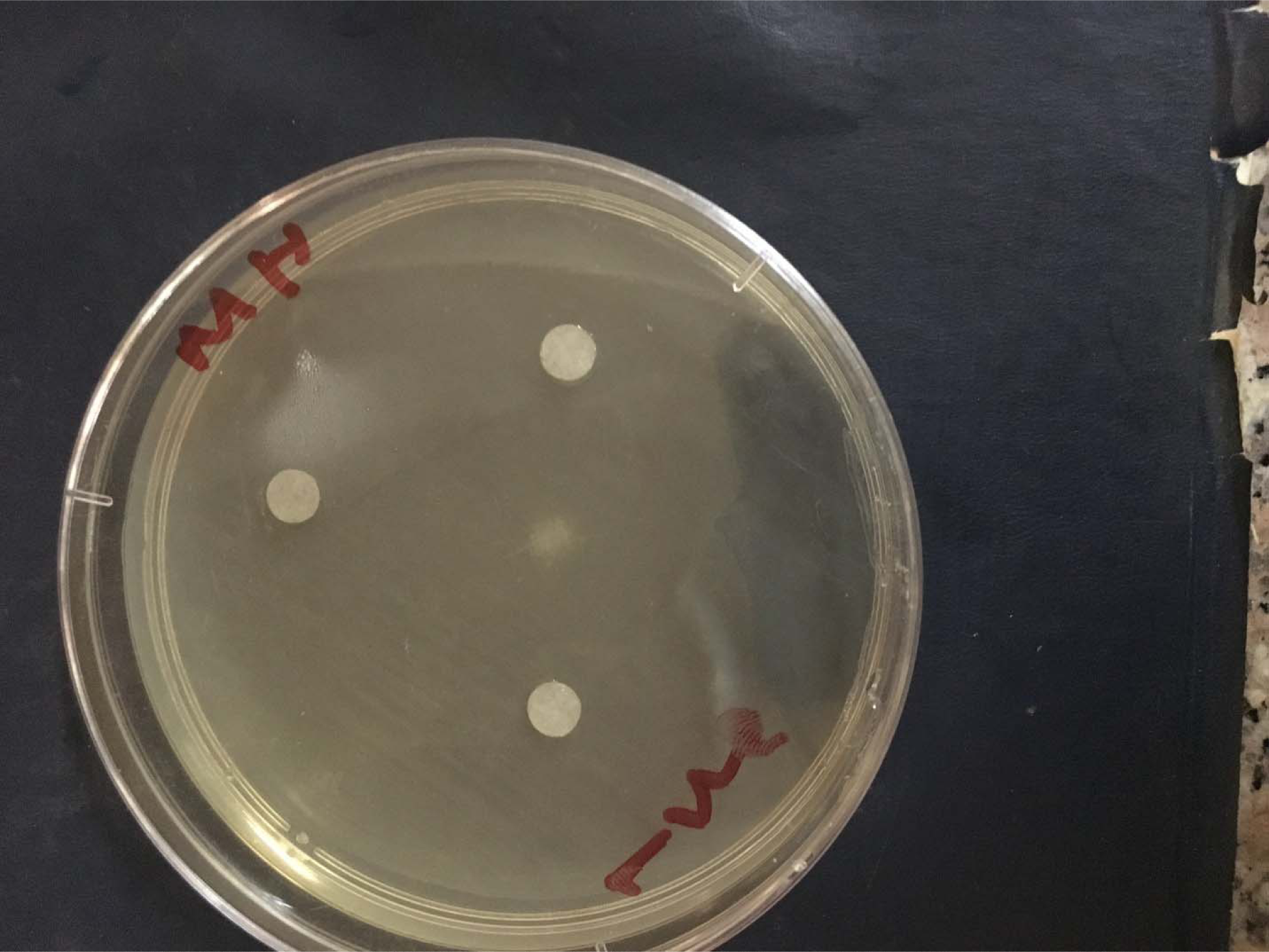
Inhibition zones around discs saturated with Ethanol 20% on a plate inoculated with E. faecalis.

The data were analyzed using SPSS 20 software, and the Least Significant Difference (LSD) test with a significance level of ≤ 0.05 was used to compare spray dried Gum Arabic, mechanically ground Gum Arabic, Chlorhexidine, and Sodium Hypochlorite.

Spray-dried Gum Arabic displayed significantly greater inhibition zone diameter (14.67 mm) against E. faecalis than mechanically ground (8.67mm) (P-value =0.02).

Gum Arabic, with both processing methods, exhibited lower antibacterial activity against E. faecalis than chlorhexidine (0.2%).However, the result was significant with the mechanically ground variant (P-value = 0.005).

Spray-dried Gum Arabic displayed a greater diameter of the inhibition zone (14.67mm) than 1% Sodium hypochlorite (12mm), but the result was statistical not significant.

Chlorhexidine (0.2%) showed a greater inhibition zone diameter (16 mm) against E. faecalis than sodium hypochlorite (1%) (12mm). However, the result was statistically insignificant.

Only Ciprofloxacin 5mcg, Levofloxacin 5mcg, Cotrimoxazole 25mcg & Tetracycline 300mcg of antibiotic list in the multidisc for Gram-positive bacteria exhibited antibacterial activity against E. faecalis (Table 2).

One sample t-test was used to compare Gum Arabic to antibiotic multidisc. The significance level was adjusted to 0.03 based on Bonferroni criteria.

Gum Arabic, with both processing methods, displayed significantly lower antibacterial activity to E. faecalis than (Ciprofloxacin 5mcg) & (Levofloxacin 5mcg) (P-values for spray-dried=0.004 / mechanically ground=0.001).

Spray-dried Gum Arabic (14.6 mm) showed almost the same antibacterial effect against E. faecalis as Cotrimoxazole 25 mcg (15mm), but the result was statistically not significant. However, mechanically ground Gum Arabic was significantly lower than Cotrimoxazole (P-value=0.003).

Spray-dried Gum Arabic displayed significantly higher antibacterial activity against E. faecalis than Tetracycline 300mcg (P-value=0.005). In contrast, mechanically ground showed lower antibacterial activity than Tetracycline with no statistical significance.

### Antibacterial susceptibility tests for serial dilutions of Gum Arabic (100, 50, 25, and 12.5 mg/ml) against Enterococcus faecalis

The antibacterial activity of spray dried GA exceeded that of mechanically ground GA in all the investigated concentrations, except for the concentration 12.5mg/ml they were similar. (Table 3) (Figures 8, 9)

**Table 3:**
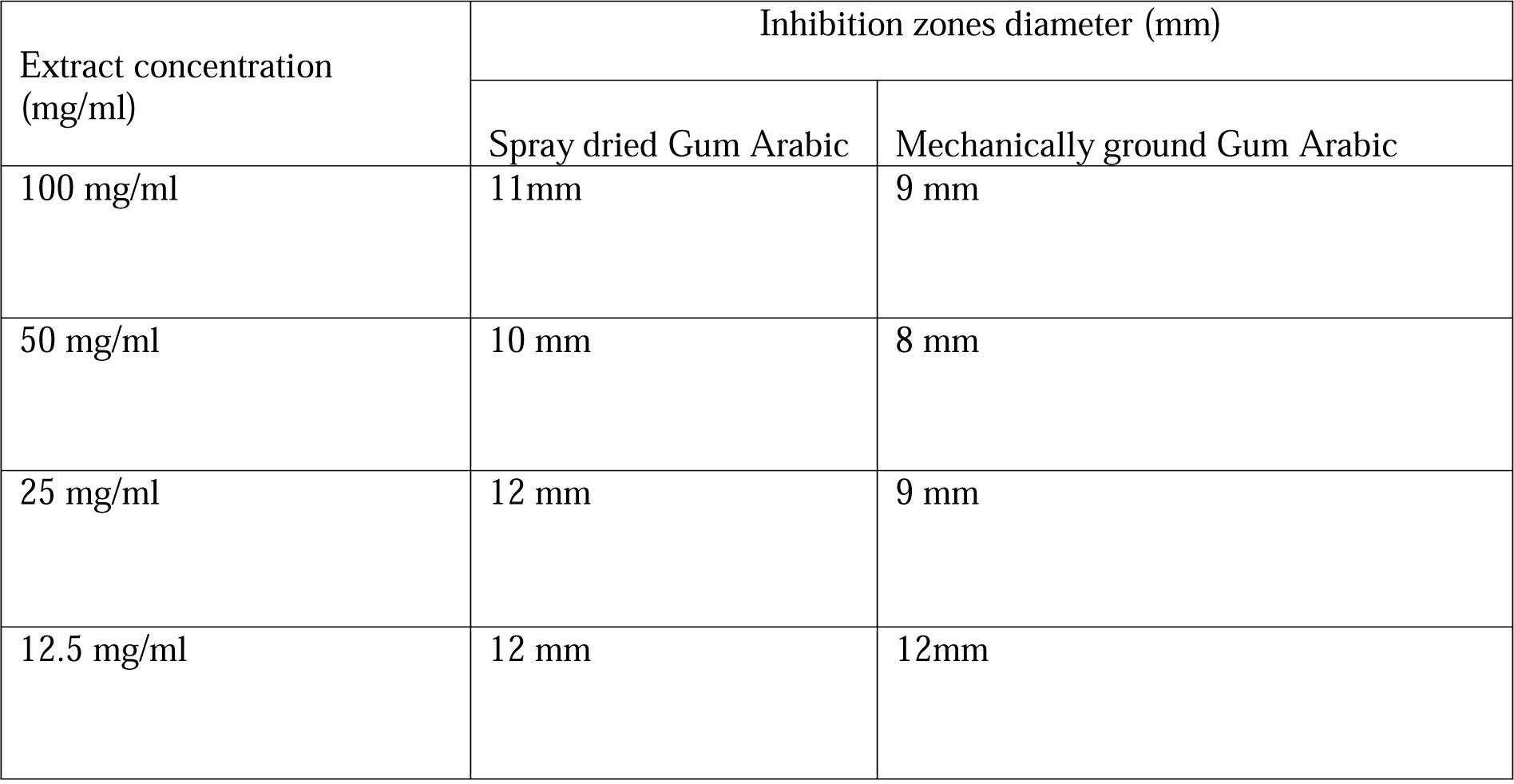
Diameter of inhibition zones produced by serial dilutions of spray-dried and mechanically ground Gum Arabic against E. faecalis.

| Extract concentration<br>(mg/ml) | Inhibition zones diameter (mm) |  |
| --- | --- | --- |
|  | Spray dried Gum Arabic | Mechanically ground Gum Arabic |
| 100 mg/ml | 11mm | 9 mm |
| 50 mg/ml | 10 mm | 8 mm |
| 25 mg/ml | 12 mm | 9 mm |
| 12.5 mg/ml | 12 mm | 12mm |

**Figure 8:**
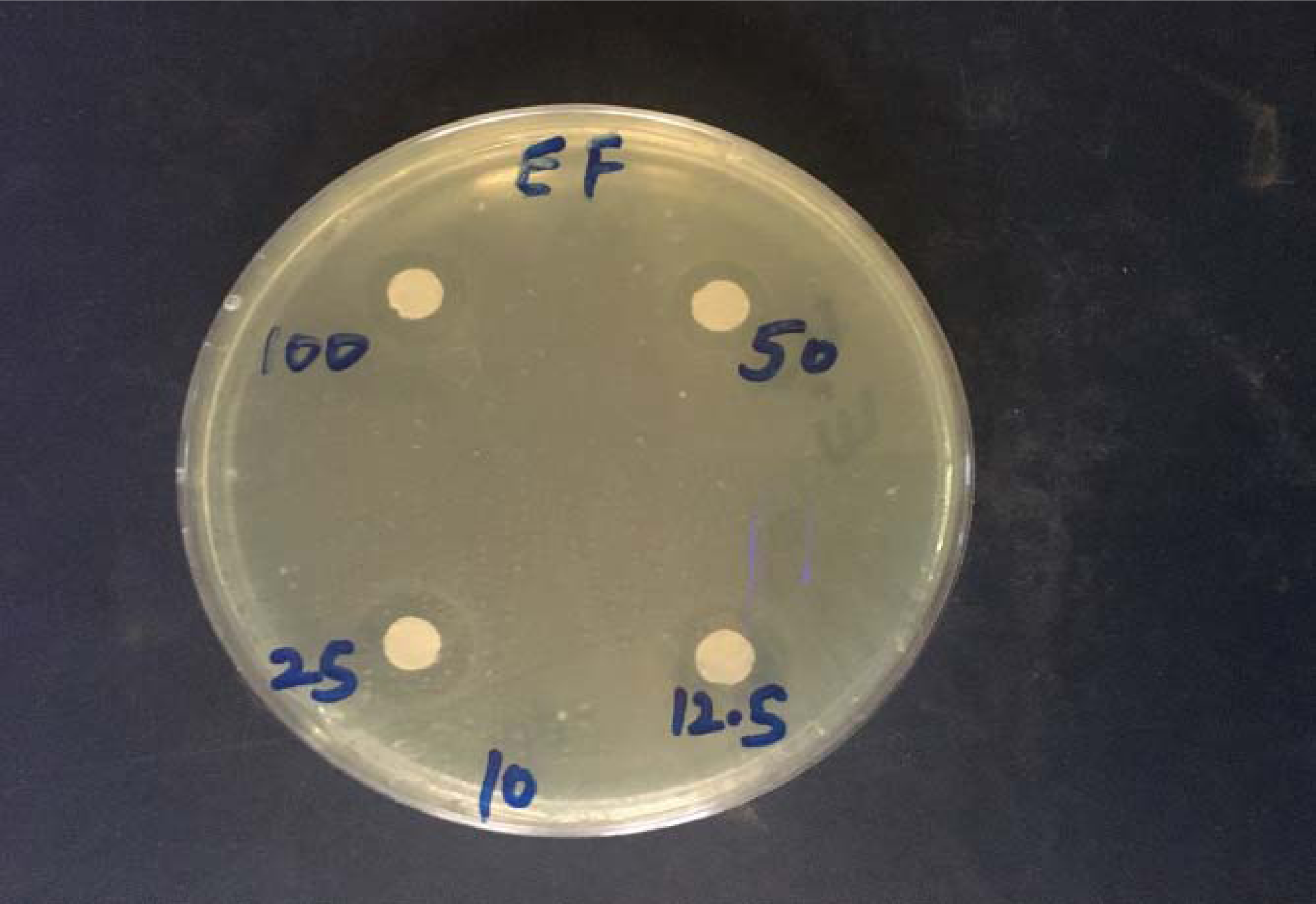
Inhibition zones around discs saturated with serial dilutions (100, 50, 25, 12.5mg/ml) of spray dried Gum Arabic on a plate inoculated with E. faecalis.

**Figure 9:**
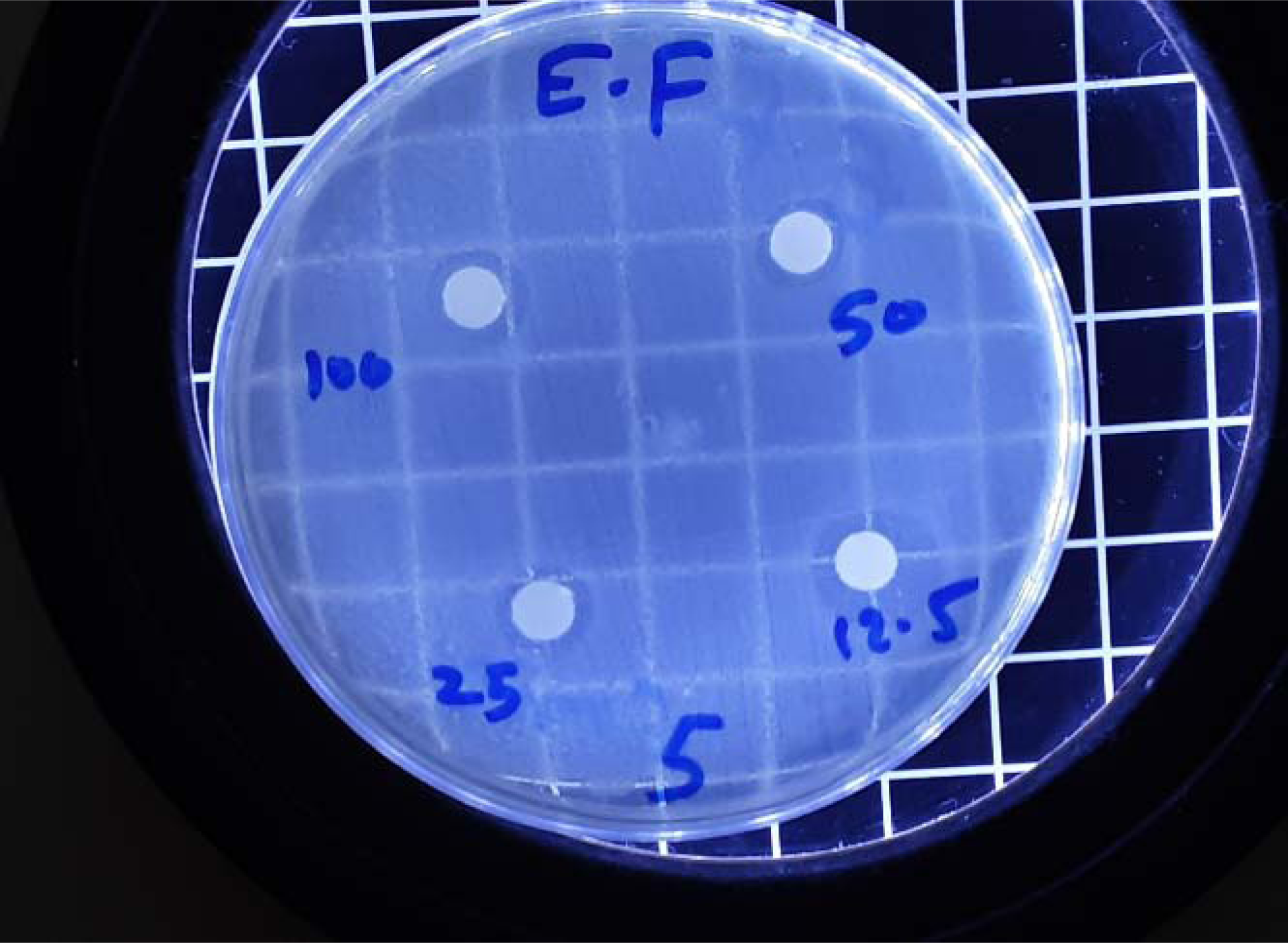
Inhibition zones around discs saturated with serial dilutions (100, 50, 25, 12.5mg/ml) of mechanically ground Gum Arabic on a plate inoculated with E. faecalis.

An Excel blot for log concentrations of Gum Arabic versus the square radii inhibition zones showed a regression line for each type of Gum Arabic (Table 4) (Figures 10, 11).

**Table 4:**
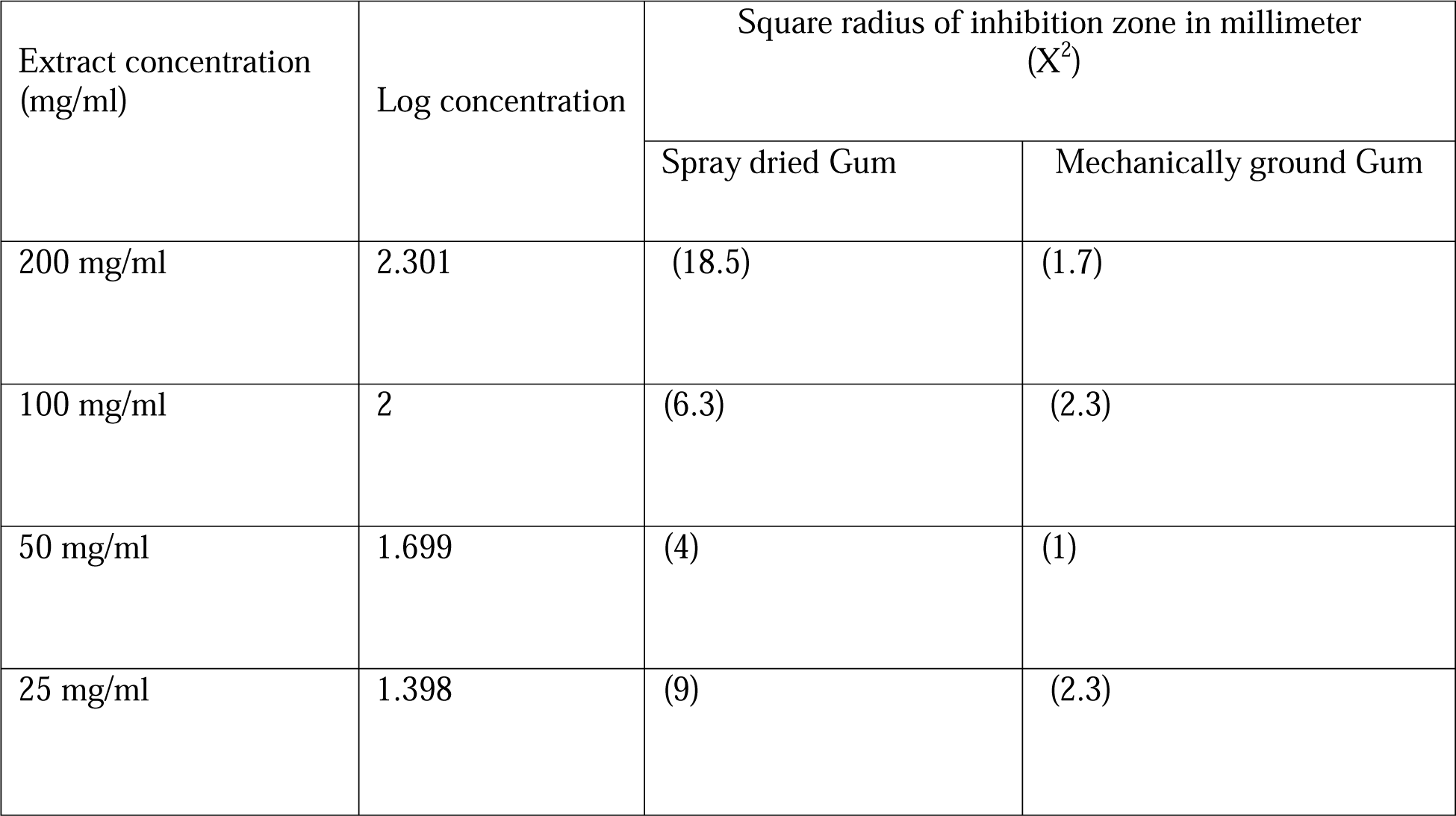
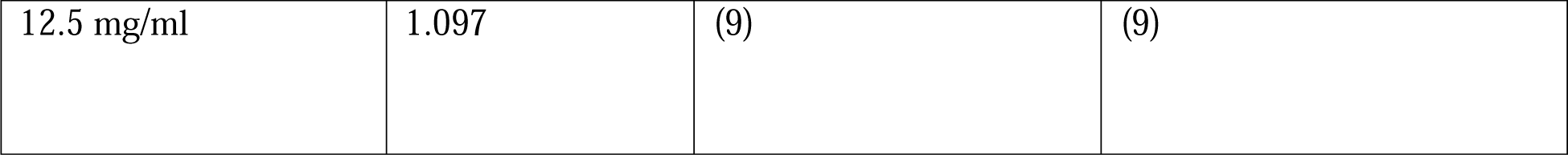
Antibacterial activity of Gum Arabic against E. faecalis represented by square radius of inhibition zone diameter versus log concentration of the extract.

| Extract concentration<br>(mg/ml) | Log concentration | Square radius of inhibition zone in millimeter<br>(X<sup>2</sup>) |  |
| --- | --- | --- | --- |
|  |  | Spray dried Gum | Mechanically ground Gum |
| 200 mg/ml | 2.301 | (18.5) | (1.7) |
| 100 mg/ml | 2 | (6.3) | (2.3) |
| 50 mg/ml | 1.699 | (4) | (1) |
| 25 mg/ml | 1.398 | (9) | (2.3) |
| 12.5 mg/ml | 1.097 | (9) | (9) |

**Figure 10:**
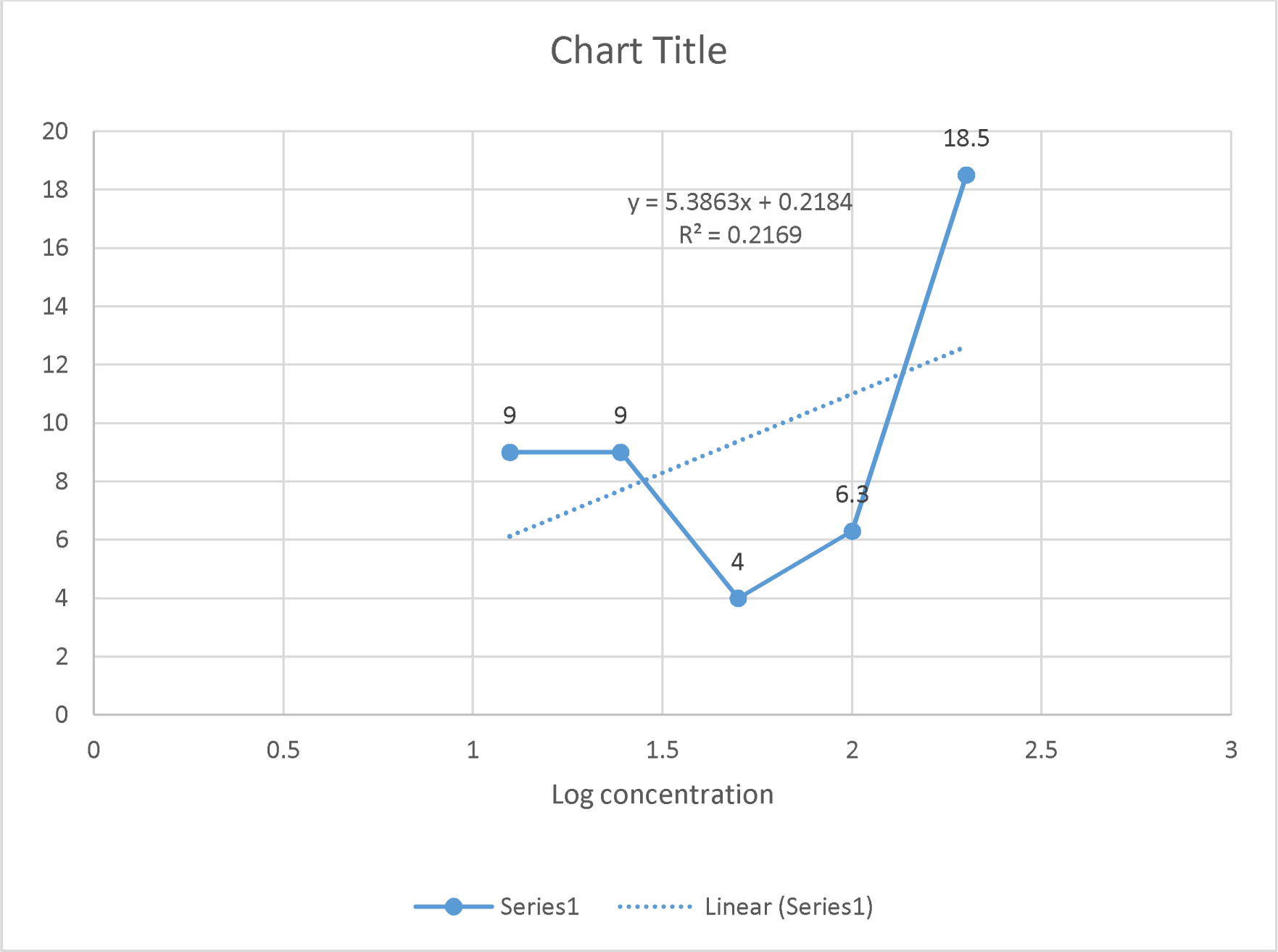
Excel plot shows the relation between log concentrations of spray-dried GA (X-axis) and square radii of inhibition zones (Y-axis) using simple regression analysis.

**Figure 11:**
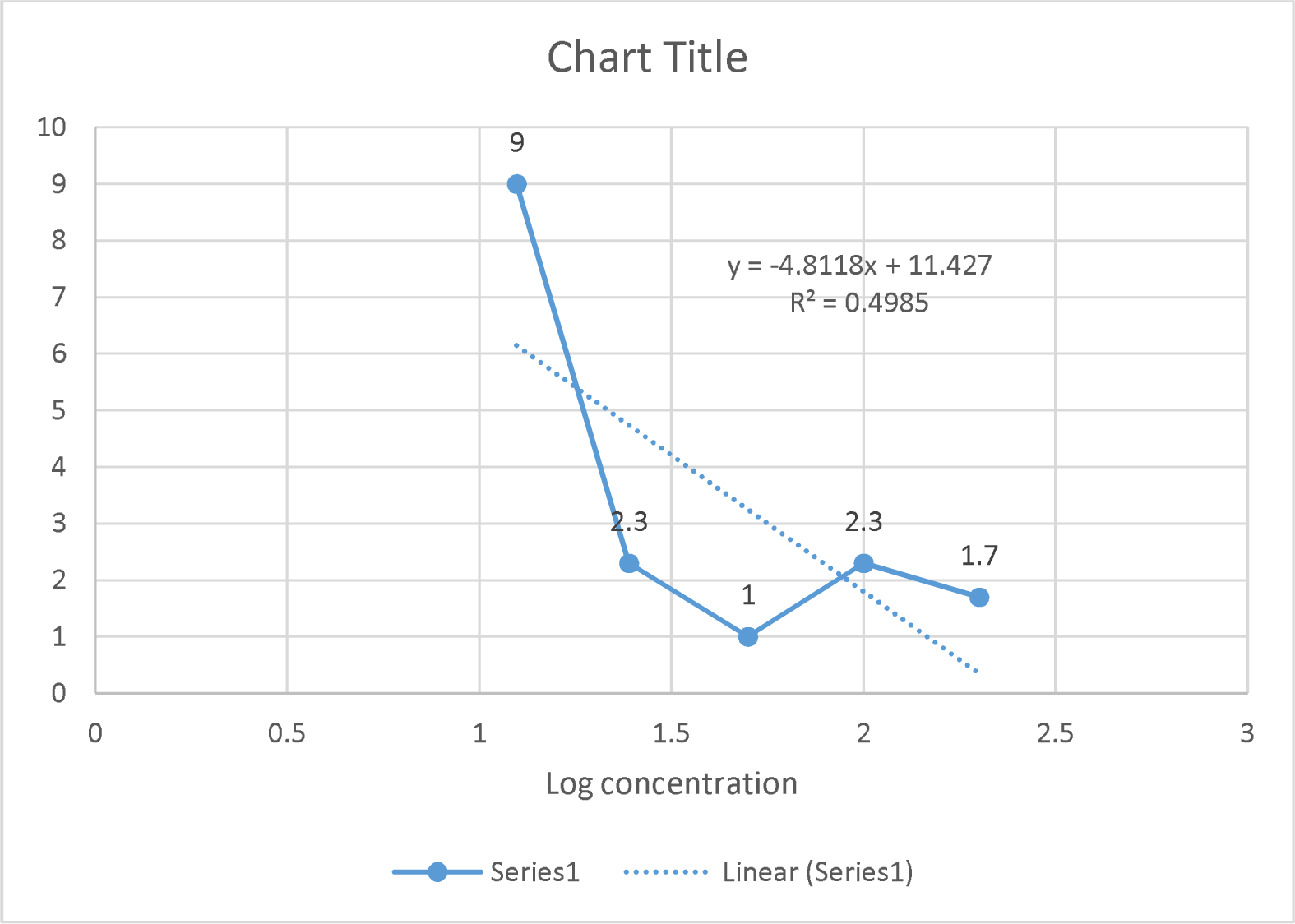
Excel plot shows the relation between log concentrations of mechanically ground GA (X-axis) and square radii of inhibition zones (Y-axis) using simple regression analysis.

The blot in Figure 10 displayed a positive relation between the concentration of spray-dried GA and inhibition. The regression line equation for spray-dried GA indicated that the inhibition zone would be zero when the log concentration of extract is -0.04 (which is the X-intercept of the line). Therefore, by calculating the antilog for -0.04, the concentration of spray-dried GA with no inhibitory effect against E. faecalis was determined as 0.9mg/ml.

In the blot shown in Figure 11, there is a negative relation between the concentration of mechanically ground GA and inhibition. The regression line equation revealed no inhibitory effect of mechanically ground GA on E. faecalis when the log concentration of extract is 2.37. As a result, the concentration was determined by calculating the antilog of 2.37, which is 234.4 mg/ml.

## Discussion

The study examined the quality of Gum Arabic processed using mechanical grinding and spray drying techniques. One of the mechanically ground samples was found to be contaminated with E. coli, whereas the other mechanically ground sample (manually) and the spray-dried sample met the Microbiological Standard for Gum Arabic. However, the spray-dried Gum Arabic showed better results than the manually ground sample. This finding supports what has been mentioned by Montenegro et al. (10) that Gum Arabic samples produced by spray drying have an advantage over raw GA as they are free of microbial contamination. The authors recommended performing microbiological analysis on Gum Arabic intended for research or medical use to ensure that only uncontaminated samples are used. comparing the amount of extract obtained from spray-dried and mechanically ground GA using 70% and 80% ethanol for extraction, displayed that the yield percentage of extract of spray-dried GA was higher than that of mechanically ground GA. This result is consistent with what was reported in the literature (19) that smaller particle size leads to a higher yield percentage. However, it was found that spray-dried GA did not produce any extract when absolute ethanol was used, while mechanically ground GA produced an extract.

In the present study, when 200 mg ethanolic extract was investigated, the spray-dried GA was found to be active antibacterial against Enterococcus faecalis. In contrast, the mechanically ground GA was found to be inactive antibacterial. The activity level was determined using Alves et al.(20), and Mukhtar & Ghori (21) scale, which categorized products as inactive if the inhibition zone was less than 9mm, partially active if it was between 9-12mm, active if it was between 13-18mm, and very active if it was more than 18mm.

There are several reasons that may explain the difference in the antimicrobial properties between the two types of Gum Arabic. One of these reasons is the variation in the particles size. The size of Gum Arabic particles varies depending on the processing method used. For instance, mechanical grinding creates particles that can reach a size of 400 μm (19), whereas spray drying produces particles of 50-100 μm (10). Larger particles take a long time to complete the extraction process (22). Lowering the particle size of the plant powder increases surface contact between plant particles and the extraction solvent to produce efficient extraction and higher quality extract. Therefore, the spray drying technique with the uniform small particle size might produce higher-quality extract with more active ingredients, leading to more antibacterial properties than mechanically ground Gum Arabic.

Another reason is the drawbacks of agar disc diffusion technique itself. One limitation of this technique is that not all materials have the same diffusing characteristics within agar. Although the same amounts of extract solutions were applied to Whatman filter paper discs, it was impossible to quantify the material diffused through the agar medium (23). Based on this drawback, the small-sized particles of spray-dried Gum Arabic may exhibit greater diffusing features in Agar compared to the larger particles of mechanically ground Gum Arabic, resulting in a larger inhibition zone gained by spray-dried Gum Arabic.

Moreover, the degree of Solubility and viscosity of Gum Arabic may considered as another reason. The molecules of Gum Arabic have a highly branched structure, which results in a compact, small hydrodynamic volume. This structure leads to the formation of a viscous solution when the concentration is high (10). The high viscosity of the solution impedes its diffusion through agar. Therefore, when GA is present at high concentrations (200mg/ml), along with large particle size from mechanical grinding, it can hinder its diffusion through agar, leading to a smaller inhibition zone diameter than spray-dried GA. This explanation is supported by the observation that mechanically ground GA achieved a similar diameter of inhibition zone as spray-dried GA at a lower concentration serial dilutions (12.5 mg/ml).

Based on the regression line equation, there is a positive relation between the log concentration of spray-dried GA and the square radius of inhibition. This implies that the inhibitory effect of spray-dried GA on E. faecalis decreases as the concentration of Gum Arabic decreases, until it reaches a concentration of 0.9 mg/ml where there is no inhibition. However, only 21.6% of the change in the square radius of inhibition can be explained by the change in concentration of spray-dried GA (R^2^=0.2169). On the other hand, the regression equation of mechanically ground GA showed a negative relation. This means that the inhibitory effect of mechanically ground GA on E. faecalis increases as the concentration of Gum Arabic decreases. In this case, 49.8% of the change in the square radius of inhibition can be explained by the change in concentration of spray-dried GA (R2 = 0.4985).

It is important to note that the current study has some limitations. The study relied on mathematical calculations based on laboratory measurements rather than complete laboratory measurements to determine the relation of Gum Arabic concentration and its inhibitory effect. Moreover, only single discs were utilized to represent the concentrations (100, 50, 25, 12.5 mg/ml) rather than using triple discs. The drawbacks of Agar Disc Diffusion method can also be added to the limitations of this study. To obtain more conclusive results regarding the relation between Gum Arabic concentration and its inhibitory effect against E. faecalis, the authors recommended using a different method than the agar disc diffusion with a wider range of concentrations, starting from above 200mg/ml and going down to below 12.5mg/ml.

Based on the variation between the antibacterial properties of spray dried and mechanically ground GA in the current study, the authors suggest that for research purposes, a chemically well-characterized preparation of Gum Arabic should be used to achieve consistency and standardization across different studies. This suggestion is in agreement with what was recommended by Ali and his colleagues (24).

## Conclusion

When extracting Gum Arabic using 70% and 80% ethanol, it was found that spray-dried GA produced more ethanolic extract than mechanically ground GA.

The method used to process Gum Arabic affects its antibacterial potency. In high concentrations, spray-dried GA is considered an active antibacterial agent against E. faecalis, while mechanically ground GA is considered non-active. When the concentration of mechanically ground GA is decreased, its inhibitory effect against E. faecalis increases, but the opposite effect is observed when using spray-dried Gum Arabic.

The antibacterial activity of spray-dried GA against E. faecalis exceeds that of Tetracycline 300mcg.

